# Genome-wide bisulfite sequencing data of normal and *Ascosphaera apis*-infected larval guts of eastern honeybee

**DOI:** 10.1101/2020.04.01.020040

**Authors:** Yu Du, Zhiwei Zhu, Huazhi Chen, Yuanchan Fan, Jie Wang, Xiaoxue Fan, Haibin Jiang, Cuiling Xiong, Yanzhen Zheng, Dafu Chen, Rui Guo

## Abstract

*Apis cerana cerana*, a subspecies of eastern honey, *Apis cerana*, plays a specific role in beekeeping industry and ecosystem in China and other Asian countries. Larvae of *A. c. cerana* can be infected by *Ascosphaera apis*, the fungal pathogen of chalkbrood. In this article, normal 4-, 5-, and 6-day-old larval guts (AcCK1, AcCK2, AcCK3) and *A. apis*-infected 4-, 5- and 6-day-old larval guts (AcT1, AcT2, AcT3) of *A. c. cerana* workers were respectively harvested followed by DNA isolation, bisulfite conversion, cDNA library construction and Illumina sequencing. Based on genome-wide bisulfite sequencing, 79167210, 82175052, 79331489, 81051009, 74742842 and 74849091 raw reads were generated from AcCK1, AcCK2, AcCK3, AcT1, AcT2 and AcT3, and after quality control, 73417030 (92.73%), 76660370 (93.27%), 71804727 (90.44%), 75046507 (92.82%), 67487782 (90.30%) and 67367023 (90.04%) clean reads were obtained, respectively. Additionally, 73333333, 76533333, 71466667, 75066667, 67590965 and 67200000 clean reads were mapped to the reference genome of *A. cerana*, including 54656767, 58583415, 54127407, 57943220, 52547867 and 51295824 unique mapped clean reads, and 8624392, 8789458, 7531333, 7747337, 6249679 and 5394174 multiple mapped clean reads. The genome-wide bisulfite sequencing data reported here can be used for genome-wide identification of 5mC methylation sites in eastern honeybee larval guts and systematic investigation of DNA methylation-mediated host response to *A. apis* infection.

**Value of the data:**

- The current dataset contributes to genome-wide identification of 5mC methylation sites in normal and *A. apis*-infected larval guts of eastern honeybee.
- The reported data could be used for systematic investigation of DNA methylation-mediated response of eastern honeybee larvae to *A. apis* infection.
- Our data offers a valuable genetic resource for better understanding epigenetic regulation mechanism involved in eastern honeybee larvae-*A. apis* interaction.

## 1. Data description

The data reported here were obtained from genome-wide bisulfite sequencing of normal and *Ascosphaera apis*-infected larval guts of *Apis cerana cerana*. The ratio of G and C among total bases derived from genome-wide bisulfite sequencing of normal 4-, 5-, and 6-day-old larval guts (AcCK1, AcCK2, AcCK3) and *A. apis*-infected 4-, 5-, and 6-day-old larval guts (AcT1, AcT2, AcT3) was obviously much lower than that of A and T (**Figure 1, Figure 2, Figure 3, Figure 4, Figure 5, Figure 6**). In total, 79167210, 82175052, 79331489, 81051009, 74742842 and 74849091 raw reads were yielded from AcCK1, AcCK2, AcCK3, AcT1, AcT2 and AcT3 groups, amounting to about 11.88 Gb, 12.33 Gb, 11.90 Gb, 12.16 Gb, 11.21 Gb and 11.23 Gb, respectively (**Table 1**). After quality control, 73417030 (92.73%), 76660370 (93.27%), 71804727 (90.44%), 75046507 (92.82%), 67487782 (90.30%) and 67367023 (90.04%) clean reads were gained, amounting to about 10.99 Gb, 11.48 Gb, 10.75 Gb, 11.24 Gb, 10.11 Gb and 10.09 Gb, respectively (**Table 1**). Additionally, the GC contents of clean reads from the aforementioned six groups were 17.33%, 17.33%, 17.83%, 17.67%, 18.17% and 19%, respectively (**Table 1**). Furthermore, 73333333, 76533333, 71466667, 75066667, 67590965 and 67200000 clean reads were mapped to the reference genome of *Apis cerana*, including 54656767, 58583415, 54127407, 57943220, 52547867 and 51295824 unique mapped clean reads, and 8624392, 8789458, 7531333, 7747337, 6249679 and 5394174 multiple mapped clean reads (**Table 2**).

**Table 1.**
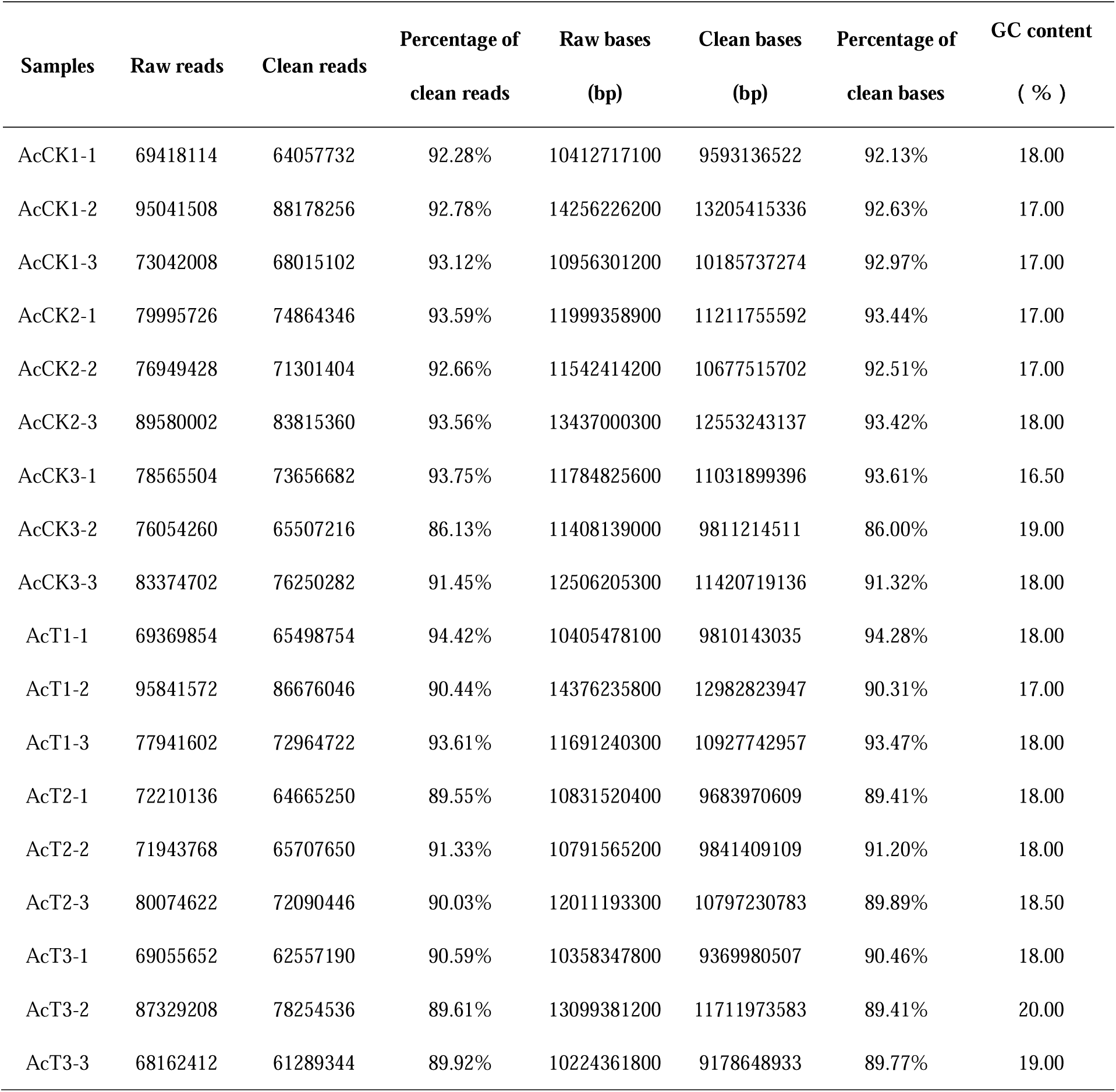
Summary of genome-wide bisulfite sequencing of *A. c. cerana* larval guts in control and treatment groups.

**Table 2.** Summary of mapping of clean reads to the reference genome of *A. cerana*.

| Samples | Clean reads | Unique mapped reads | Multiple mapped reads | Unmapped reads | Mapping ratio |
| --- | --- | --- | --- | --- | --- |
| AcCK1-1 | 64000000 | 48299599 | 7244955 | 8455446 | 86.79% |
| AcCK1-2 | 88000000 | 63877814 | 11025730 | 13096456 | 85.12% |
| AcCK1-3 | 68000000 | 51792887 | 7602491 | 8604622 | 87.35% |
| AcCK2-1 | 74800000 | 56188178 | 9396741 | 9215081 | 87.68% |
| AcCK2-2 | 71200000 | 53830285 | 8624498 | 8745217 | 87.72% |
| AcCK2-3 | 83600000 | 65731781 | 8347135 | 9521084 | 88.61% |
| AcCK3-1 | 73200000 | 55699500 | 9611064 | 7889436 | 89.22% |
| AcCK3-2 | 65200000 | 50506793 | 5305352 | 9387855 | 85.60% |
| AcCK3-3 | 76000000 | 56175929 | 7677583 | 12146488 | 84.02% |
| AcT1-1 | 66000000 | 50787912 | 6594927 | 8617161 | 86.94% |
| AcT1-2 | 86400000 | 65727414 | 9537471 | 11135115 | 87.11% |
| AcT1-3 | 72800000 | 57314334 | 7109613 | 8376053 | 88.49% |
| AcT2-1 | 64665248 | 49607260 | 6509890 | 8548098 | 86.78% |
| AcT2-2 | 66107648 | 50574533 | 6014641 | 9518474 | 85.60% |
| AcT2-3 | 72000000 | 57461807 | 6224507 | 8313686 | 88.45% |
| AcT3-1 | 62400000 | 49484583 | 5664547 | 7250870 | 88.38% |
| AcT3-2 | 78000000 | 56976725 | 5996043 | 15027232 | 80.73% |
| AcT3-3 | 61200000 | 47426164 | 4521931 | 9251905 | 84.88% |

**Figure 1.**
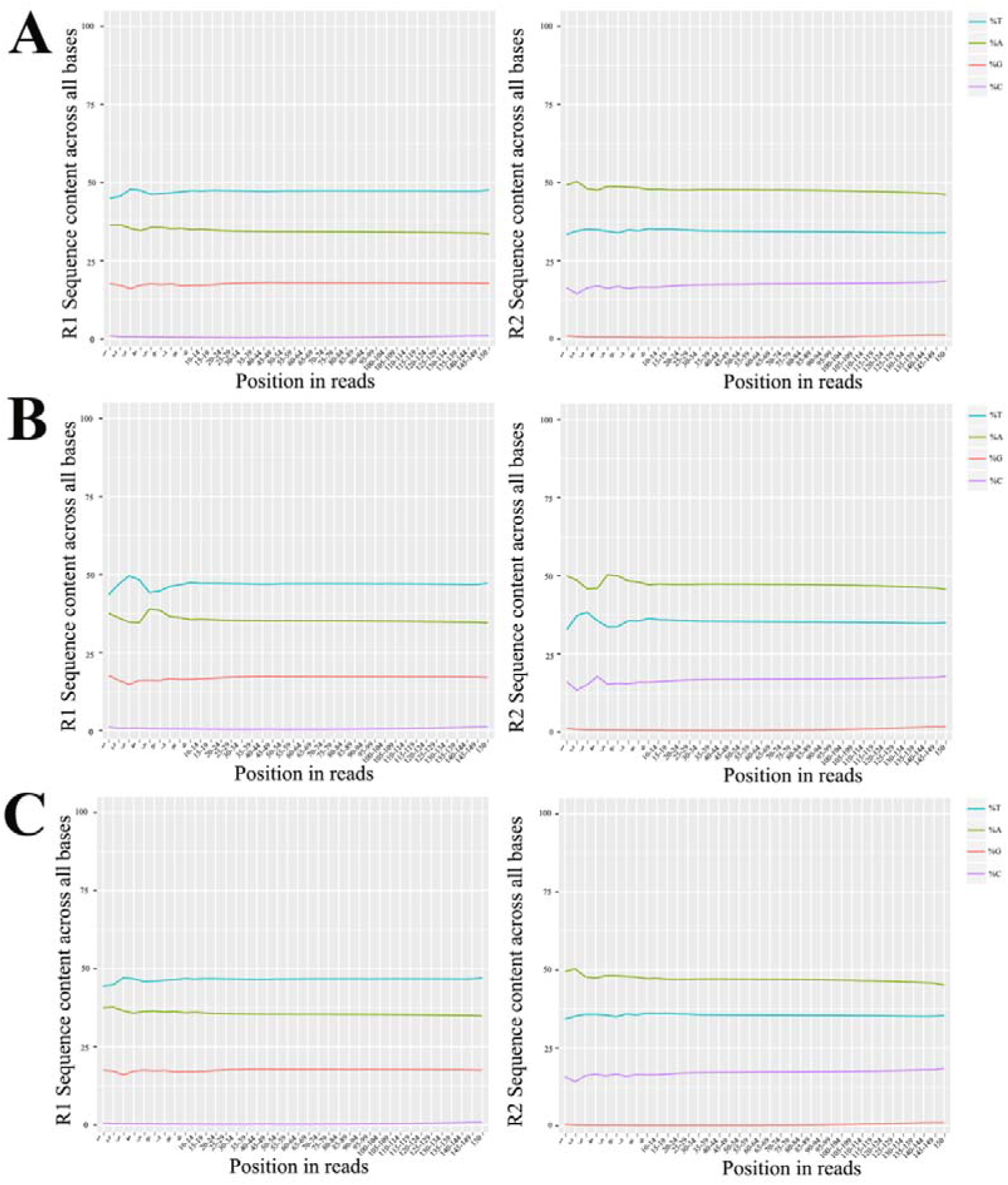
Ratio of four different bases from normal 4-day-old larval gut of *A. c. cerana*. The *x* axis indicates the position in reads, and the *y* axis indicates the ratio of a single base among total bases.

**Figure 2.**
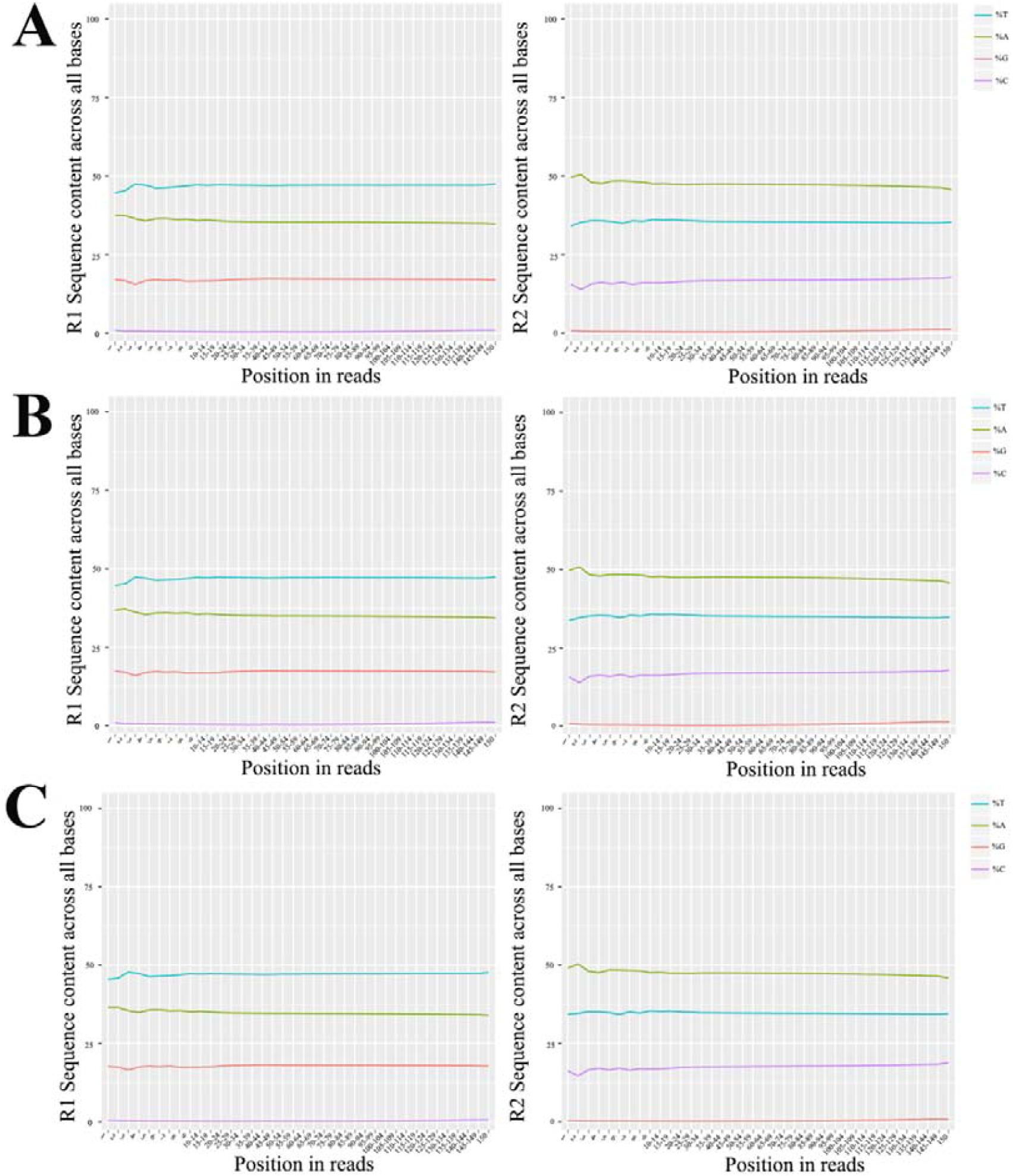
Ratio of four different bases from normal 5-day-old larval gut of *A. c. cerana*. The *x* axis indicates the position in reads, and the *y* axis indicates the ratio of a single base among total bases.

**Figure 3.**
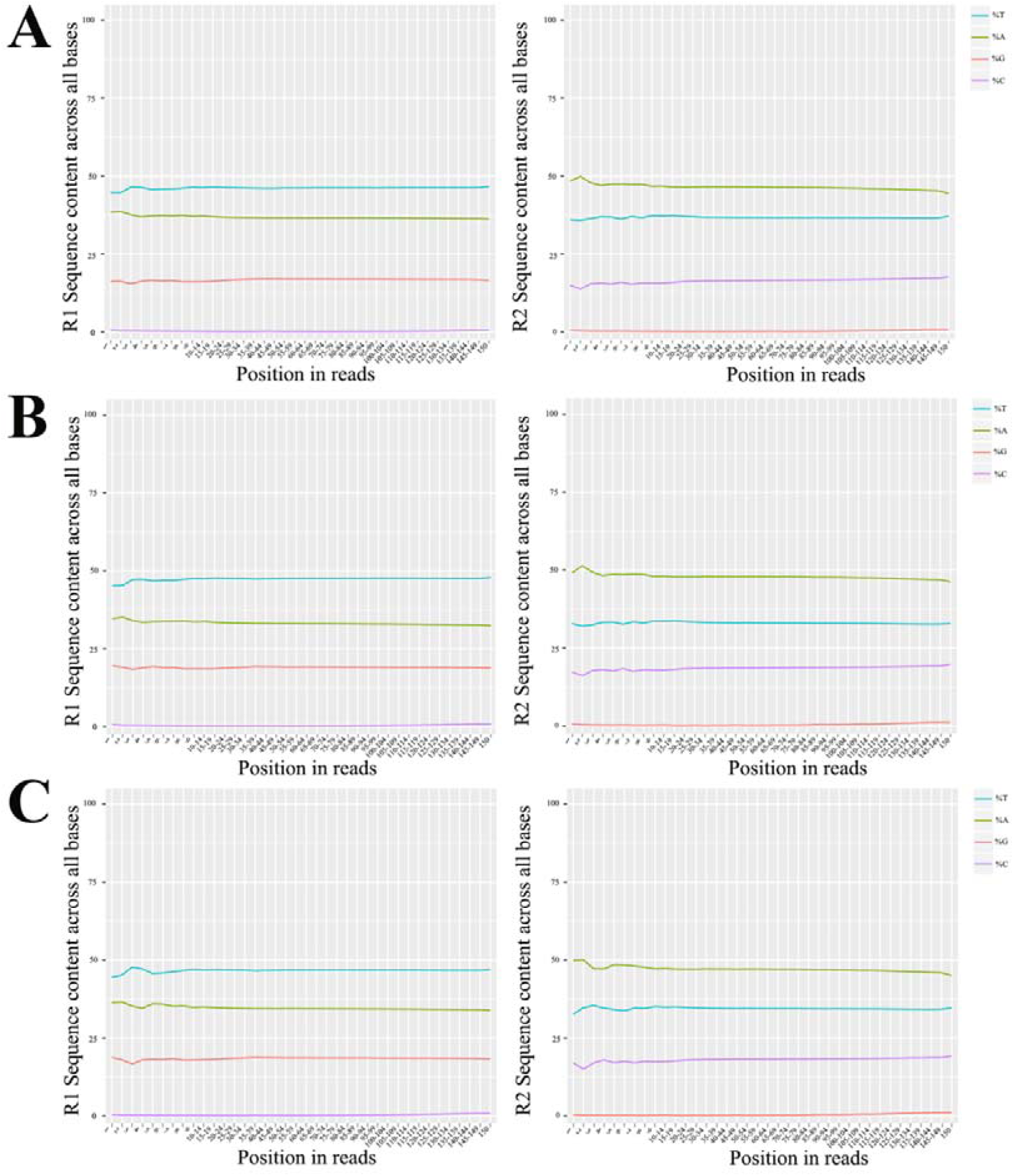
Ratio of four different bases from normal 6-day-old larval gut of *A. c. cerana*. The *x* axis indicates the position in reads, and the *y* axis indicates the ratio of a single base among total bases.

**Figure 4.**
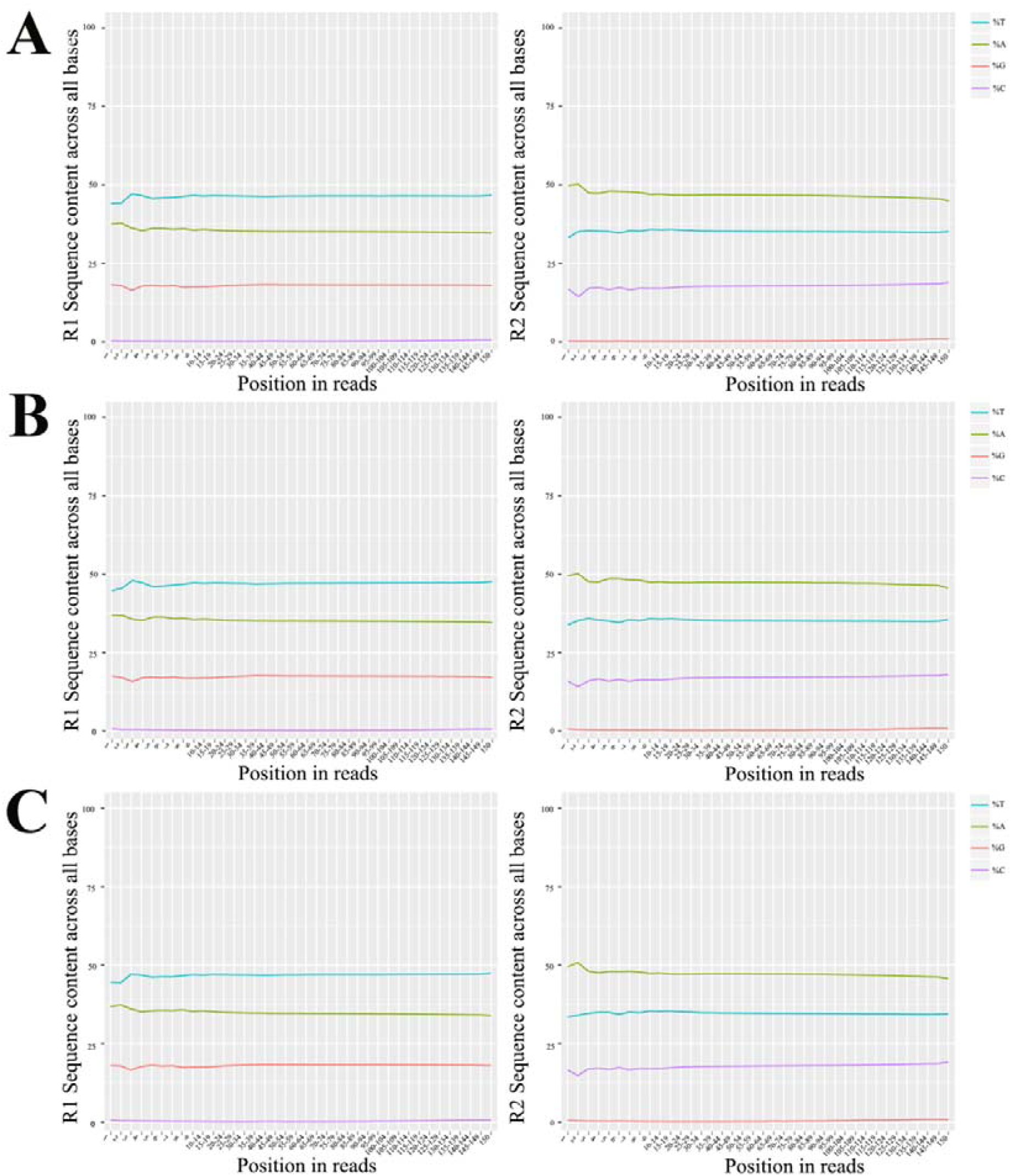
Ratio of four different bases from *A. apis*-infected 4-day-old larval gut of *A. c. cerana*. The *x* axis indicates the position in reads, and the *y* axis indicates the ratio of a single base among total bases.

**Figure 5.**
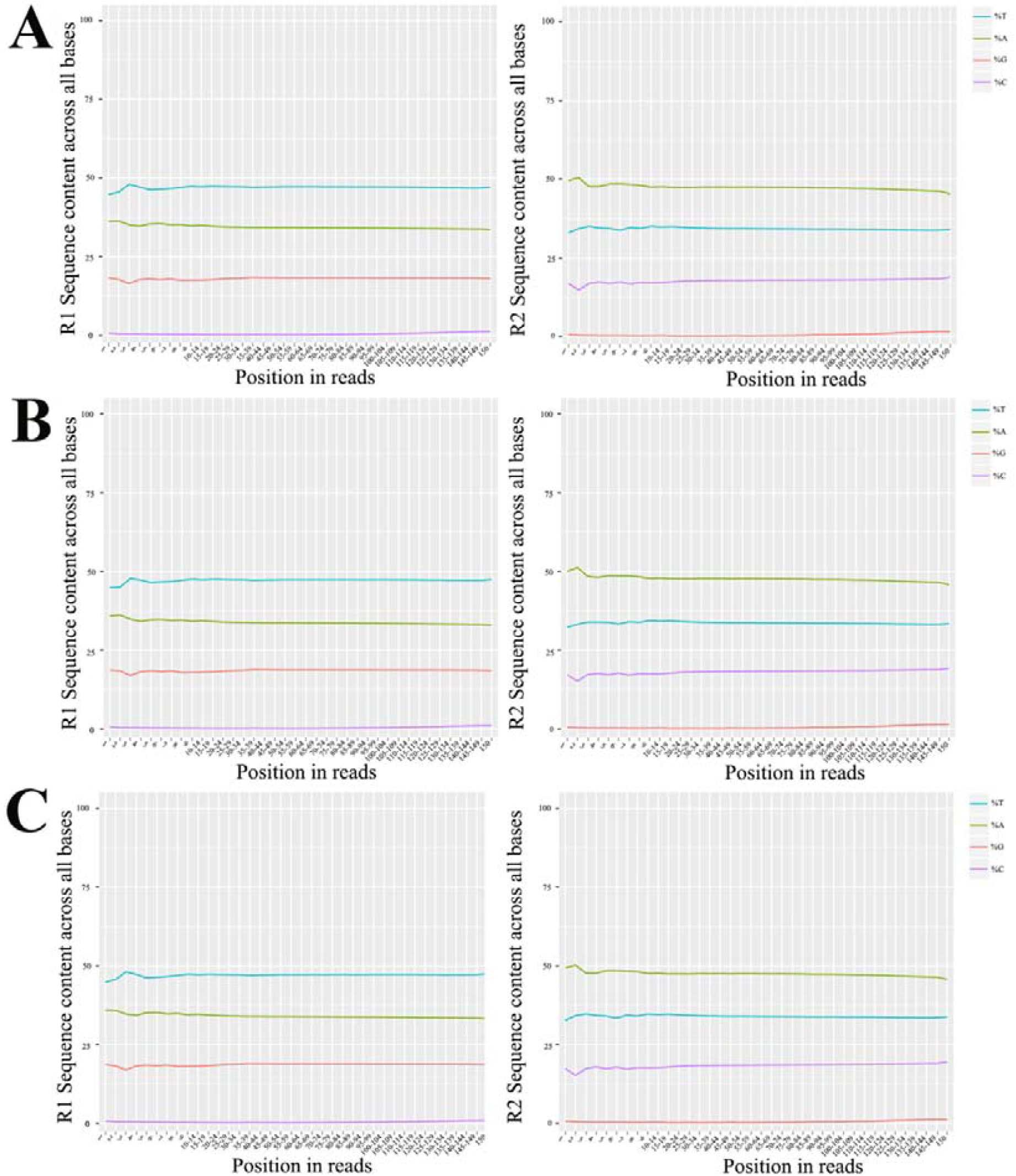
Ratio of four different bases from *A. apis*-infected 5-day-old larval gut of *A. c. cerana*. The *x* axis indicates the position in reads, and the *y* axis indicates the ratio of a single base among total bases.

**Figure 6.**
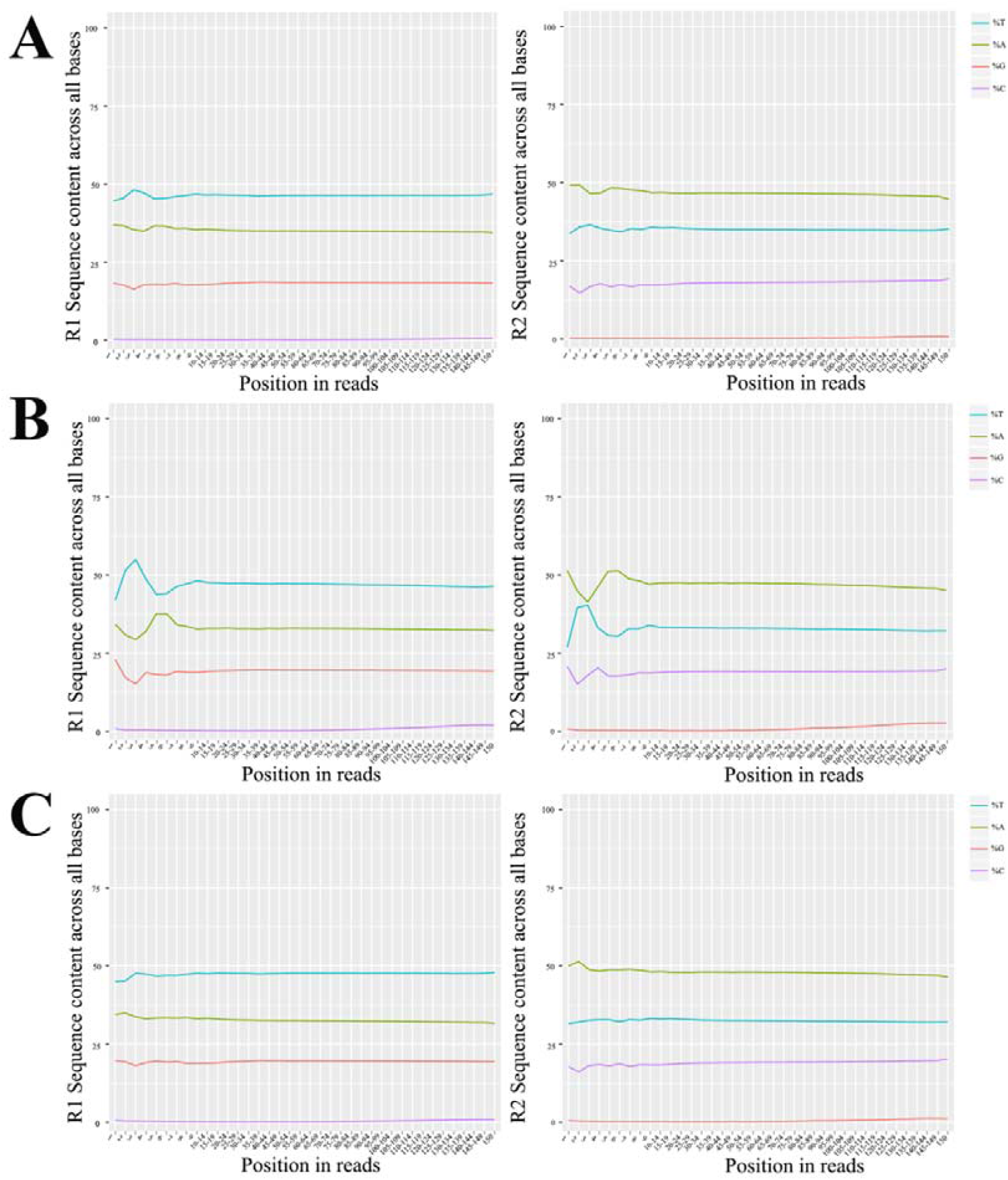
Ratio of four different bases from *A. apis*-infected 6-day-old larval gut of *A. c. cerana*. The *x* axis indicates the position in reads, and the *y* axis indicates the ratio of a single base among total bases.

## 2. Experimental Design, Materials, and Methods

### 2.1. Preparation of eastern honeybee larval gut samples

Two-day-old larvae were removed from the combs with a Chinese grafting tool and seriously transferred to a droplet of 10 μL diet. Larvae were feed once a day with 20 μL (3-day-old), 30 μL (4-day-old), 40 μL (5-day-old) and 50 μL (6-day-old) diet, 150 μL diet total per larva. Honeybee larvae were reared in 48-cell culture plates according to the method described by Peng et al. [1].

*A. apis* [2] was previously isolated and conserved at the Honeybee Protection Laboratory of the College of Animal Sciences (College of Bee Science) in Fujian Agriculture and Forestry University. Before experimental inoculation, *A. apis* was cultured at 33±0.5 °C on plates of Potato dextrose agar (PDA) medium followed by purification of fungal spores using the method developed by Jensen et al. [3].

Three-day-old larvae in treatment groups were reared with artificial diet containing *A. apis* spores (1×10^7^ spores/mL), while 3-day-old larvae in control groups were reared with artificial diet without fungal spores. Then, 4-, 5- and 6-day-old larvae in both control groups and treatment groups were reared with artificial diet without *A. apis* spores. Using the method previously developed in our lab [4], guts of 4-, 5- or 6-day-old larvae from *A. apis*-treated groups (n=21) and normal groups (n=21) were respectively harvested.

### 2.2. Bisulfite conversion and library preparation

Firstly, genomic DNA was respectively extracted from AcCK1, AcCK2, AcCK3, AcT1, AcT2, and AcT3 groups using a Universal Genomic DNA Extraction Kit (TaKaRa, Tokyo, Japan) according to the manufacturer’s instruction. Next, DNA was bisulfite treated using a Zymo Research EZ DNA methylaiton-Glod Kits (Zymo Research, Irvine, CA, USA). DNA concentration and integrity were examined by a NanoDrop 2000 spectrophotometer (Thermo Fisher Scientific, Waltham, MA, USA) and agarose gel electrophoresis, respectively. Finally, the eighteen libraries were constructed with TruSeq DNA Methylation Kit (Illumina, San Diego, CA, USA) following the manufacturer’s protocol.

### 2.3. Next-generation sequencing and data processing

The constructed eighteen cDNA libraries were sequenced on an Illumina HiSeq X Ten platform by OE Biotech Co., Ltd. (Shanghai, China) and 150 bp paired-end reads were generated. After being processed with fastp [5], clean reads were gained by removing reads containing adapter, reads containing ploy-N and low quality reads from raw reads, and used for downstream analyses. Further, the clean reads were aligned to the *A. cerana* genome (assembly ACSNU-2.0) using the genome-wide bisulfite sequencing mapping program (BSMAP, v2.74) [6]. Mapping ratio was then calculated for each sample group.

## Acknowledgments

This research was financially supported by the Earmarked Fund for China Agriculture Research System (No. CARS-44-KXJ7), the Science and Technology Planning Project of Fujian Province (No. 2018J05042), the Teaching and Scientific Research Fund of Education Department of Fujian Province (No. JAT170158), the Outstanding Scientific Research Manpower Fund of Fujian Agriculture and Forestry University (No. xjq201814), and the Scientific and Technical Innovation Fund of Fujian Agriculture and Forestry University (No. CXZX2017342, No. CXZX2017343).

